# Characterizing glucokinase variant mechanisms using a multiplexed abundance assay

**DOI:** 10.1101/2023.05.24.542036

**Authors:** Sarah Gersing, Thea K. Schulze, Matteo Cagiada, Amelie Stein, Frederick P. Roth, Kresten Lindorff-Larsen, Rasmus Hartmann-Petersen

**Affiliations:** The Linderstrøm-Lang Centre for Protein Science, Department of Biology, University of Copenhagen, Ole Maaløes Vej 5, DK-2200 Copenhagen, Denmark; Donnelly Centre, University of Toronto, Toronto, ON, M5S 3E1, Canada; Department of Molecular Genetics, University of Toronto, Toronto, ON, M5S 1A8, Canada; Lunenfeld-Tanenbaum Research Institute, Sinai Health, Toronto, ON, M5G 1X5, Canada; Department of Computer Science, University of Toronto, Toronto, ON, M5T 3A1, Canada

## Abstract

Amino acid substitutions can perturb protein activity in multiple ways. Understanding their mechanistic basis may pinpoint how residues contribute to protein function. Here, we characterize the mechanisms of human glucokinase (GCK) variants, building on our previous comprehensive study on GCK variant activity. We assayed the abundance of 95% of GCK missense and nonsense variants, and found that 43% of hypoactive variants have a decreased cellular abundance. By combining our abundance scores with predictions of protein thermodynamic stability, we identify residues important for GCK metabolic stability and conformational dynamics. These residues could be targeted to modulate GCK activity, and thereby affect glucose homeostasis.

## Introduction

Protein function is crucial for cellular and organismal homeostasis, but can be perturbed by missense variants through various mechanisms. For instance, amino acid substitutions in active site residues can directly affect protein activity, but in general such residues only constitute a small fraction of a protein. Conversely, many residues affect the thermodynamic folding stability of a protein. As most proteins need to fold into the native conformation to be functional, a widespread mechanism for missense variants is to decrease protein stability, leading to protein unfolding, degradation and a decreased protein abundance in the cell (***Yue et al., 2005***; ***Sahni et al., 2015***; ***Nielsen et al., 2017***; ***Scheller et al., 2019***; ***Abildgaard et al., 2019***). In addition, variants may affect other functional sites than catalytic residues, such as interaction interfaces and allosteric sites. Missense variants may therefore result in the same phenotype through multiple independent mechanisms. Understanding the molecular mechanisms of protein variants not only improves our general understanding of protein function but is also important for interpreting and interfering with the effects of disease-causing variants.

Pathogenic variants in the glucokinase gene (*GCK*) are linked to at least three diseases. Heterozygous variants that increase activity lead to hyperinsulinemic hypoglycemia (HH, MIM# 601820), where insulin is secreted at low blood glucose levels (***Christesen et al., 2002***; ***Glaser et al., 1998***). Conversely, variants that decrease activity are linked to diabetes; GCK-maturity-onset diabetes of the young (GCK-MODY, MIM# 125851) when heterozygous (***Froguel et al., 1992***; ***Hattersley et al., 1992***) and permanent neonatal diabetes mellitus (PNDM, MIM# 606176) if homozygous or compound heterozygous (***Njølstad et al., 2001, 2003***). These glucose homeostasis diseases arise due to improper insulin secretion, which in pancreatic *β*-cells is regulated by the rate of glucose phosphorylation, catalyzed by GCK (***German, 1993***; ***Meglasson and Matschinsky, 1986***; ***Meglassun et al., 1984***).

GCK is a 465-residue monomeric protein that folds into a small and a large domain (***Kamata et al., 2004***). Between the two domains is the single active site where glucose binds and becomes phosphorylated to form glucose-6-phosphate. Binding of glucose to GCK modulates the enzyme’s conformational landscape which includes multiple stable conformations (***Larion et al., 2010***; ***Sternisha et al., 2020***). In absence of glucose, GCK primarily populates the inactive super-open state, characterized by a large opening angle between the two domains and intrinsical disorder of an active site loop (residues 150–179) (***Kamata et al., 2004***). Upon glucose binding, GCK shifts towards a more compact active state, known as the closed state (***Kamata et al., 2004***). Here, the distance between the two domains is reduced, the small domain is structurally re-organized, and the 150–179 loop folds into a *β*-hairpin, collectively resulting in a catalytically active conformation. The conformational dynamics between inactive and active states occur on a millisecond timescale that is comparable to *k*_*cat*_ (***Larion et al., 2015***), which enables the dynamics to modulate GCK activity. Therefore, GCK has a sigmoidal response to glucose, which is essential for appropriate GCK activity (***Kamata et al., 2004***; ***Sternisha and Miller, 2019***).

Previously, using functional complementation in yeast, we characterized the activity of 9003 out of 9280 possible (97%) GCK missense and nonsense variants (***Gersing et al., 2022***). Accordingly, we now know the functional impact of most variants. However, the mechanisms leading to altered enzyme activity remain largely unknown. Prior mechanistic analyses of a few hyperactive variants found that some altered the dynamics and/or structure of the 150–179 loop (***Whittington et al., 2015***), while others lead to a more compact conformation in absence of glucose, similar to the closed state (***Larion et al., 2012***; ***Heredia et al., 2006a***,b). Building on this, we found that a conformational shift towards the active state could be a widespread mechanism for hyperactive variants (***Gersing et al., 2022***). The mechanisms of hypoactive variants include reduced structural stability and cellular abundance (***Kesavan et al., 1997***; ***Burke et al., 1999***), which was found to be a major determinant of the phenotypic severity in PNDM patients (***Raimondo et al., 2014***). In addition, a conformational shift towards the inactive state has been predicted to be the mechanism of five hypoactive variants using molecular dynamics simulations (***Zhang et al., 2006***). However, the mechanistic basis of especially hypoactive variants remains to be examined more broadly.

Here, we use a yeast-based growth assay to determine the abundance of 8822 (95%) GCK missense and nonsense variants. Abundance was decreased by amino acid substitutions in buried regions of the large domain. As enzyme activity was also decreased in these regions, decreased abundance is likely a major mechanism for decreased activity in the large domain. Conversely, in the small domain, variants in general had little effect on abundance. Instead we found that hypoactive variants affected the conformational dynamics of GCK, as previously seen for hyperactive variants. Collectively, our results expand the knowledge on the mechanisms of GCK disease-causing variants, and illuminate the interplay between protein dynamics and abundance in determining GCK function.

## Results and Discussion

### Measuring glucokinase variant abundance using the DHFR-PCA

In order to assay GCK variant abundance, we used the Dihydrofolate Reductase Protein-Fragment Complementation Assay (DHFR-PCA) (***Pelletier et al., 1999***; ***Campbell-Valois et al., 2005***; ***Levy et al., 2014***; ***Faure et al., 2022***). In this system a methotrexate-resistant mutant of mouse DHFR is split into two fragments. One fragment (DHFR[F3]) is fused to the protein of interest, here GCK, while the other fragment (DHFR[1,2]) is over-expressed freely. Both the fusion protein and fragment are expressed in wild-type yeast cells, which are grown on media containing methotrexate to inhibit the endogenous DHFR. If the fusion protein is abundant in the cell, the two DHFR fragments will reconstitute to form functional DHFR, thus enabling the cell to grow on methotrexate medium. However, if the fusion protein has a low abundance in the cell, less of the functional DHFR will form, leading to slower growth. In this way, yeast growth reports on protein abundance (Fig. 1A).

**Figure 1.**
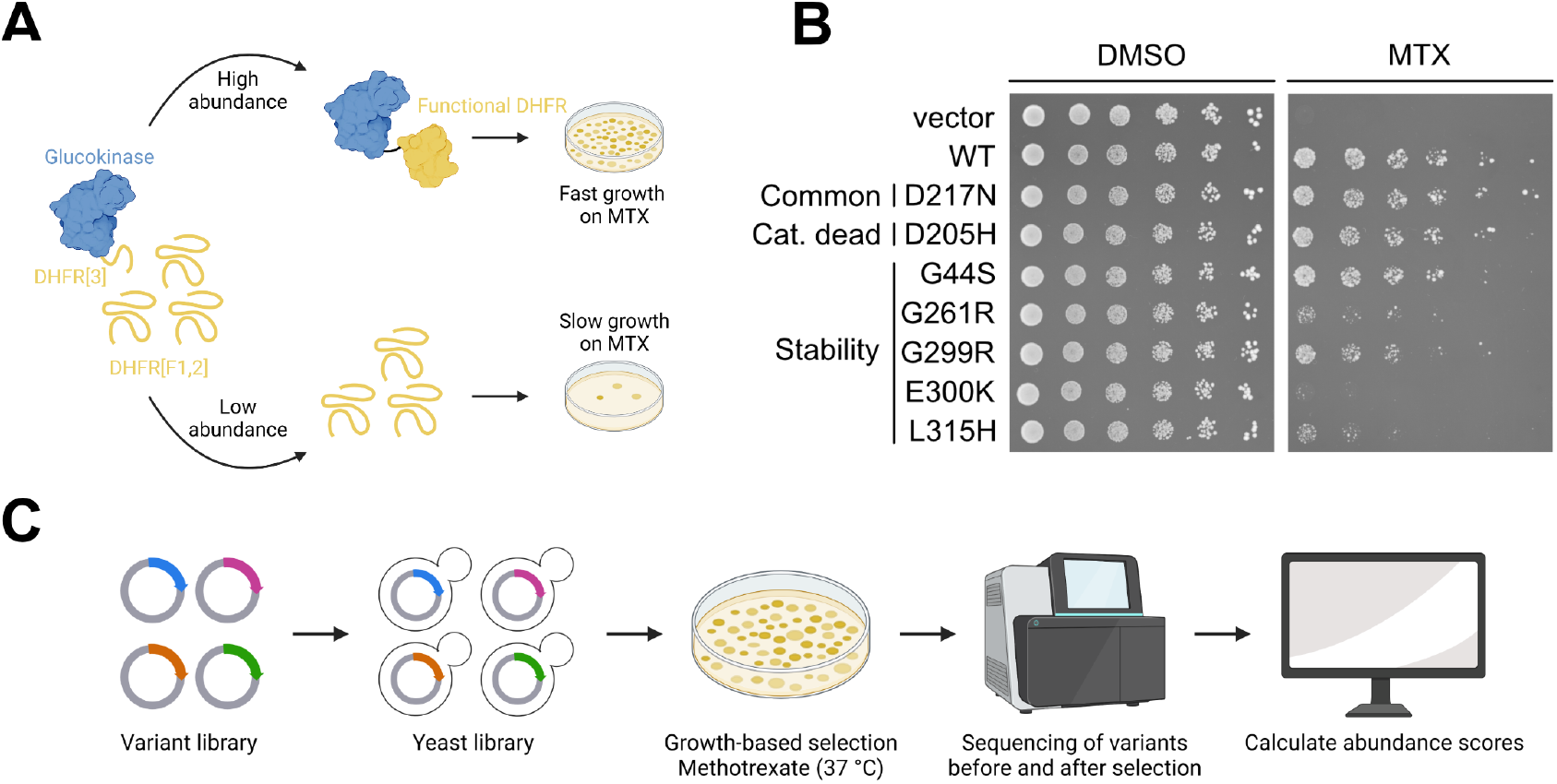
Measuring glucokinase variant abundance by DHFR-PCA. (A) Overview of the Dihydrofolate Reductase Protein-Fragment Complementation Assay (DHFR-PCA). (B) Low-throughput test of the DHFR-PCA. Selected glucokinase variants expressed in wild-type yeast cells were grown on medium without (DMSO) or with methotrexate (MTX) to assess their impact on cellular protein abundance. The vector control did not contain the DHFR-PCA sequences. (C) Overview of the multiplexed assay for glucokinase variant abundance.

To test the dynamic range of the DHFR-PCA for GCK missense variants, we assayed the wildtype protein and seven selected variants in low-throughput. Yeast expressing wild-type GCK grew on methotrexate medium, while an empty vector control showed no growth (Fig. 1B). The common variant D217N (***Karczewski et al., 2020***) and a catalytically-inactive variant D205H (***Kamata et al., 2004***; ***García-Herrero et al., 2012***) grew similar to wild-type GCK (Fig. 1B), suggesting a wild-type-like abundance as expected. The five remaining variants (G44S, G261R, G299R, E300K, and L315H) are disease-linked (***Gloyn et al., 2004***; ***Gidh-Jain et al., 1993***; ***Pruhova et al., 2010***) and were previously predicted by thermodynamic stability calculations (ΔΔ*G*) to be destabilized (***Gersing et al., 2022***). In addition, E300K is a well-studied unstable GCK variant (***Kesavan et al., 1997***; ***Burke et al., 1999***). Accordingly, all five variants showed reduced growth compared to wild-type GCK (Fig. 1B), albeit for the G44S and G299R variants this effect was less pronounced. In conclusion, the DHFR-PCA detected the low-abundance variants, and can therefore assess GCK variant abundance.

Next, we multiplexed the DHFR-PCA to widely assess the abundance of GCK missense variants (Fig. 1C). Previously, we generated a library of GCK variants (***Gersing et al., 2022***). This library was cloned into the DHFR-PCA vector to generate an abundance variant library, which was transformed into wild-type yeast. Following outgrowth, the yeast library was grown at 37 °C on methotrexate medium for four days to select for abundance. Then, variants were sequenced before and after selection, and sequencing data were analyzed to obtain abundance scores.

### A map of glucokinase variant abundance

We scored the abundance of 8822 missense and nonsense variants (95%) (Fig. 2A). As an initial quality control check of the abundance scores, we found that the variants tested in low-throughput scored as expected (Fig. S1A). In addition, the distributions of nonsense and synonymous variants were separated, while the scores of missense variants spanned from synonymous-like to nonsense-like (Fig. 2B). Consistent with expectations, nonsense mutations at most positions were not tolerated except for at the extreme C-terminus (Fig. 2A). To further validate the abundance scores, we examined the cellular protein levels of 11 GCK variants expressed with an N-terminal GFP-tag, using western blotting. The protein levels quantified from western blots correlated with abundance scores (Pearson’s r: 0.80, p-value: 2e-08) (Fig. S1BC). We note that variants with an abundance score below 0.5 all showed low cellular protein abundance, and that differences in scores below 0.5 may not translate to changes in cellular protein levels. Despite this limitation, the abundance scores reflect cellular protein abundance of the GCK variants.

**Figure 2.**
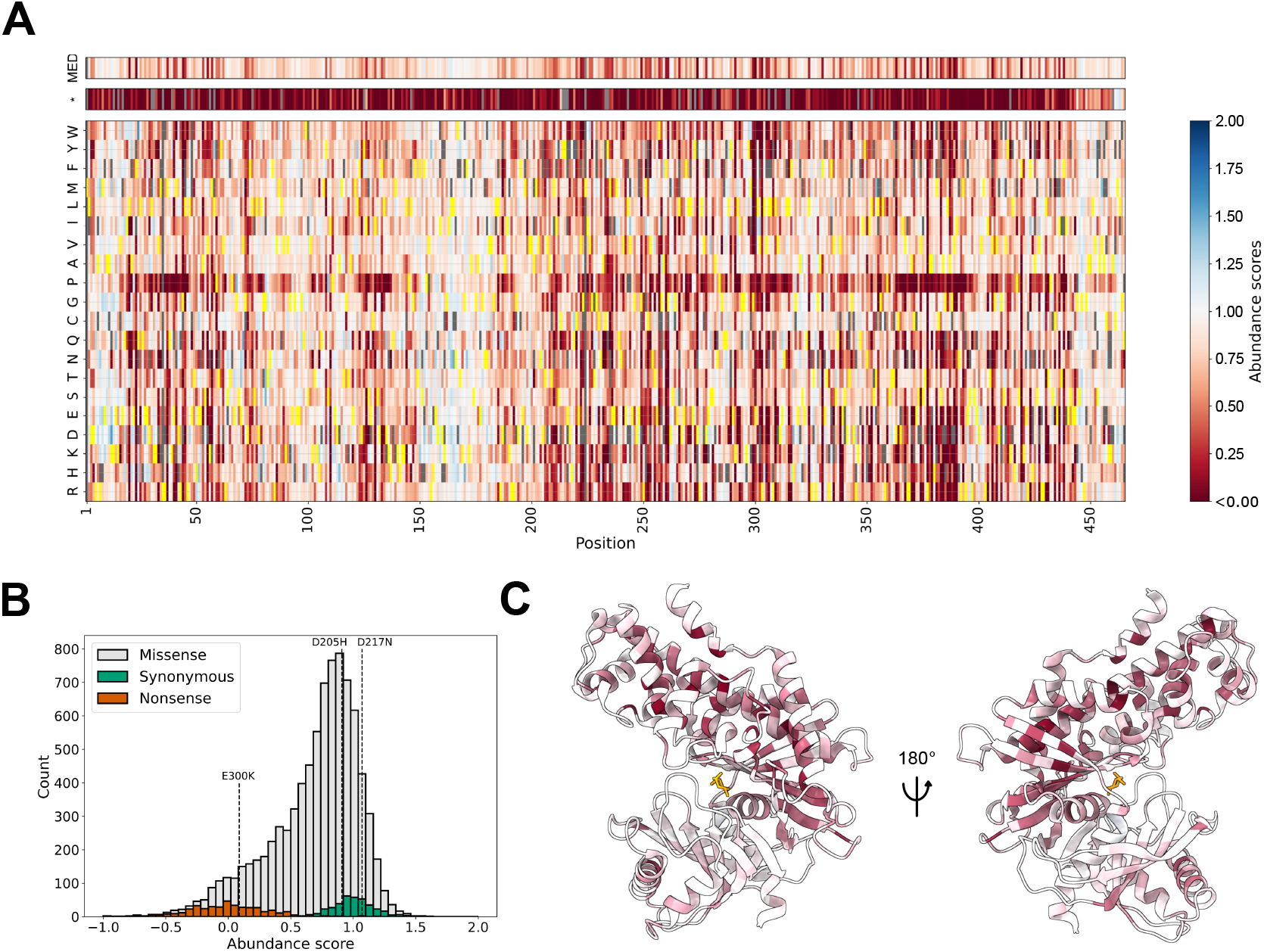
Map of glucokinase variant abundance. (A) Heatmap showing the abundance scores of 8822 missense and nonsense (*) glucokinase variants. The median score at each position is shown at the top (MED). The wild-type amino acid at each position is shown in yellow. Missing variants are shown in grey. (B) Abundance score distributions of glucokinase missense, synonymous and nonsense variants. Stippled lines indicate the scores of three variants tested in low-throughput to be unstable (E300K) or wild-type-like (D205H and D217N). (C) The closed active state of glucokinase colored by median abundance scores. The coloring scheme is the same as in panel A. Glucose is shown in orange. PDB: 1V4S.

Having validated the abundance scores, we examined variant effects structurally, mapping the median abundance score at each position onto the structure of glucose-bound GCK. This revealed that the small domain tolerated mutations at most positions (Fig. 2C), potentially due to the domain’s conformational heterogeneity and dynamic nature (***Kamata et al., 2004***; ***Larion et al., 2012***). In contrast, the large domain is more static (***Kamata et al., 2004***; ***Larion et al., 2012***). Accordingly, while surface-exposed residues seemed mutation-tolerant, most buried residues in the large domain appeared to destabilize GCK when mutated (Fig. 2C, Fig. S2), suggesting that a decrease in protein abundance is a general mechanism for loss-of-function variants in this domain.

### Mechanistic analyses of hypo- and hyperactive glucokinase variants

As decreased protein stability is a major cause of loss-of-function variants (***Yue et al., 2005***; ***Redler et al., 2016***), we next examined the relation between GCK variant abundance and activity. Previously, we generated a map of GCK variant activity (***Gersing et al., 2022***). Using these activity scores, we examined how many hypoactive variants were associated with decreased abundance using the activity and abundance scores of 9019 variants (including missense, synonymous, and nonsense variants). First, we defined an abundance score threshold below which variants are categorized as having low abundance (Fig. S3). The threshold for low activity was previously defined to be 0.66 (***Gersing et al., 2022***). Using these thresholds, a large fraction (43%) of variants with low activity also decreased abundance (Fig. 3A), in line with prior analyses (***Sahni et al., 2015***; ***Jepsen et al., 2020***; ***Cagiada et al., 2021***). These variants were enriched in the large domain, as 33% of residues in the large domain had a median abundance below 0.58, compared to 10% of small-domain residues (Fig. S4). The remaining 57% low-activity variants appeared to loose activity through other mechanisms than abundance. Conversely, 25% of the low-abundance variants scored as wild-type-like or hyperactive in the activity assay (Fig. 3A). Potentially, some of these variants might be less active at 37 °C, the temperature at which abundance was assayed. Alternatively, some variants might reduce abundance but increase specific activity, resulting in a wild-type-like or increased activity score, as the activity assay also to some extent reflects variant abundance. In conclusion, decreased abundance appears to be a major mechanism for GCK variants with decreased activity, in particular in the large domain, although the association between abundance and activity is not simple.

**Figure 3.**
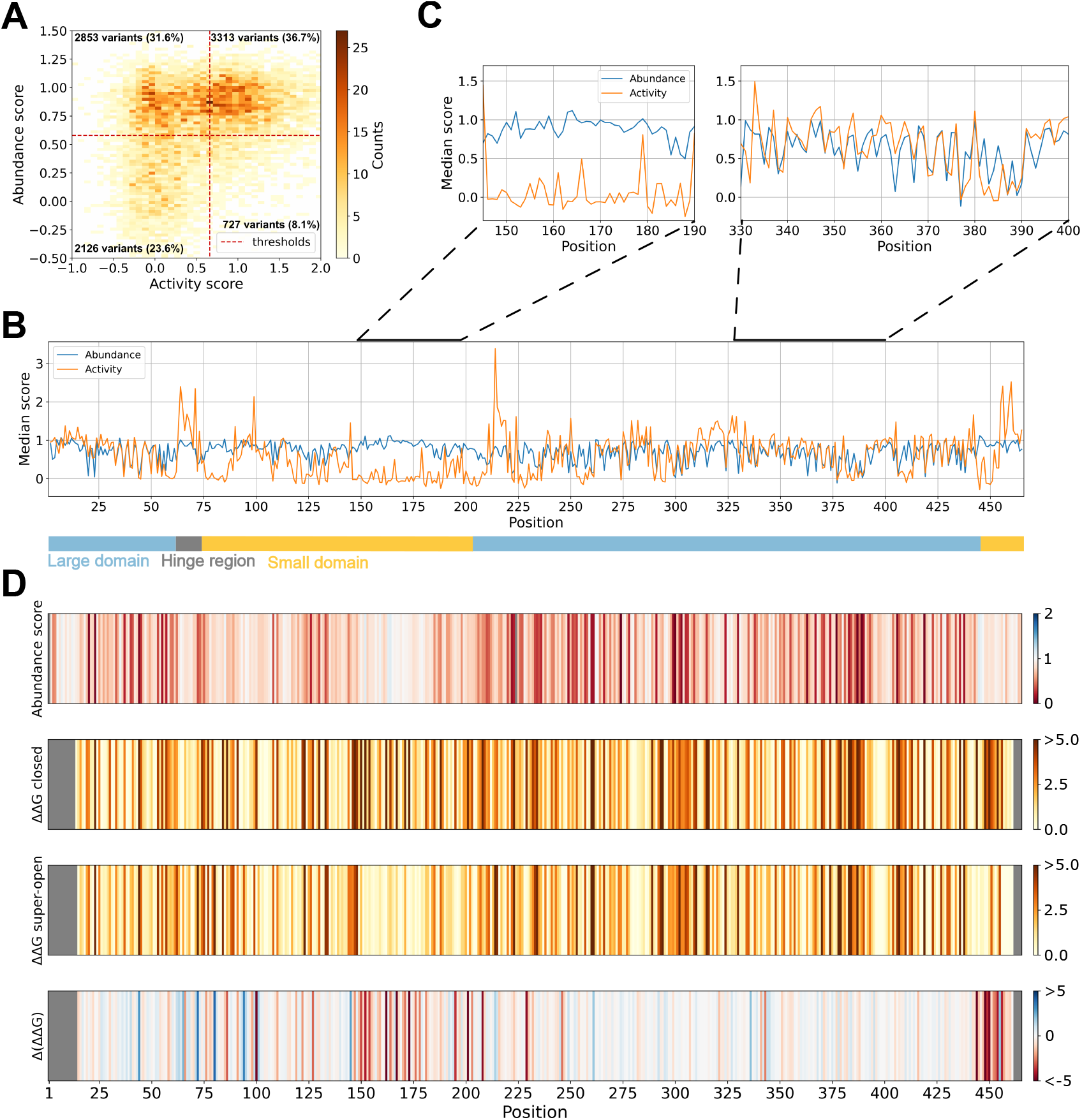
Changes in glucokinase activity explained by decreased abundance and conformational shifts. (A) Abundance and activity scores of 9019 missense, nonsense and synonymous glucokinase variants shown as a 2D histogram. The thresholds for low abundance (0.58) and low activity (0.66) are indicated as red stippled lines. The number and percentage of variants falling within each quadrant are reported. (B). The median abundance and activity of variants at each position of the glucokinase sequence is shown as a line plot. The regions forming the hinge region (grey) and the large (light blue) and small (light orange) domains are represented as a bar at the bottom. (C) Plots zooming in on regions 145–190 (left) and 330–400 (right) from panel B. (D) Barcode plots showing the median abundance score, predicted change in protein thermodynamic stability (ΔΔ*G*, kcal/mol) using the closed active state, (ΔΔ*G*, kcal/mol) using the super-open inactive state, and difference (Δ(ΔΔ*G*)) between the two ΔΔ*G* predictions. For the bottom plot, red indicates that variants at these positions are predicted to destabilize the closed state more, while at blue positions variants are predicted to destabilize the super-open state more. PDBs: 1V4S (closed) and 1V4T (super-open). The ΔΔ*G* data were obtained from (***Gersing et al., 2022***) except for the 157–179 region in the super-open state. For all panels, the data on glucokinase variant activities were obtained from (***Gersing et al., 2022***).

In order to identify the regions of GCK where changes in activity upon mutation are not explained by abundance, we compared the median activity and abundance scores along the GCK sequence (Fig. 3B). In some regions, the two medians showed a good correlation (Fig. 3B and C right panel), suggesting that loss-of-activity variants at these positions are caused by decreased abundance. In contrast, some regions showed large deviations between the two scores (Fig. 3B and C left panel). In general, regions with increased activity appeared unaffected in the abundance assay (Fig. 3B), suggesting that a changed abundance is not a common mechanism for hyperactive variants. Notably, nearly all regions where variants increased or decreased activity without affecting abundance are part of the small domain (Fig. 3B).

The small domain attains several conformations during GCK’s catalytic cycle (***Kamata et al., 2004***; ***Larion et al., 2012***). Consequently, small-domain variants might affect GCK activity by altering GCK dynamics. Such a mechanism is well-established for hyperactive variants (***Heredia et al., 2006a***,b; ***Gersing et al., 2022***; ***Larion et al., 2012***; ***Whittington et al., 2015***). For hypoactive variants, molecular dynamics simulations have predicted five small-domain variants to shift GCK towards inactive conformations (***Zhang et al., 2006***). In addition, we previously used predictions of protein thermodynamic stability (ΔΔ*G*) for the structures of super-open and closed GCK to examine a conformational shift mechanism (***Gersing et al., 2022***). Although we mostly focused on hyperactive variants, we found two regions around residues 150 and 450 where hypoactive variants were predicted to shift GCK towards the inactive state. Accordingly, the region around residue 450, corresponding to helix 13, was previously found to modulate the allosteric properties of GCK (***Larion and Miller, 2009***).

Our prior mechanistic analysis of hypoactive variants was limited by residues 157–179 missing from the crystal structure of super-open GCK. To examine this region further, we created five different structural models of the super-open state that included the 157–179 loop region, assuming that the region is disordered, as previously seen for all prominent substates of unliganded GCK (***Larion et al., 2015***). For all five models, we predicted the change in protein thermodynamic stability using Rosetta (***Park et al., 2016***), and used the average ΔΔ*G*s from the five models for the missing loop residues to supplement our previous predictions (***Gersing et al., 2022***). As previously, we calculated the difference between the ΔΔ*G*s in the closed and super-open state (ΔΔ*G*_super-open_−ΔΔ*G*_closed_). Variants with a high negative score are predicted to shift GCK towards the inactive (super-open) state, given that they do not severely destabilize the super-open conformation, which would likely lead to decreased cellular abundance. Many residues were on average predicted to shift GCK towards the super-open conformation upon mutation, and these spanned the entire 150–179 region (Fig. 3D). Variants in the 150–179 region might therefore severely decrease activity without affecting abundance by shifting GCK into an inactive state.

### Variants in the 150–179 region affect glucokinase conformational dynamics

To substantiate a conformational shift mechanism for hypoactive variants experimentally, we focused on the 150–179 region. If the region’s disorder in the super-open state results in mutational tolerance with respect to abundance, then any disordered sequence should be tolerated without perturbing GCK protein abundance. To test this, we replaced the region spanning residues 150– 179 with a GS-repeat sequence of either 30 or 6 residues (Fig. 4A). The resulting mutants retained no detectable activity (Fig. 4B), as expected, but did not affect the cellular protein level of GCK compared to wild-type (Fig. 4C). When we further examined abundance using the DHFR-PCA, the mutants grew similar to wild-type GCK (Fig. 4D), again supporting that abundance was not affected. In conclusion, the region spanning residues 150–179 can be replaced by six residues (GSGSGS) or a 30 residue GS repeat without affecting GCK cellular abundance. This is consistent with the region being highly tolerant towards mutations in the super-open state.

**Figure 4.**
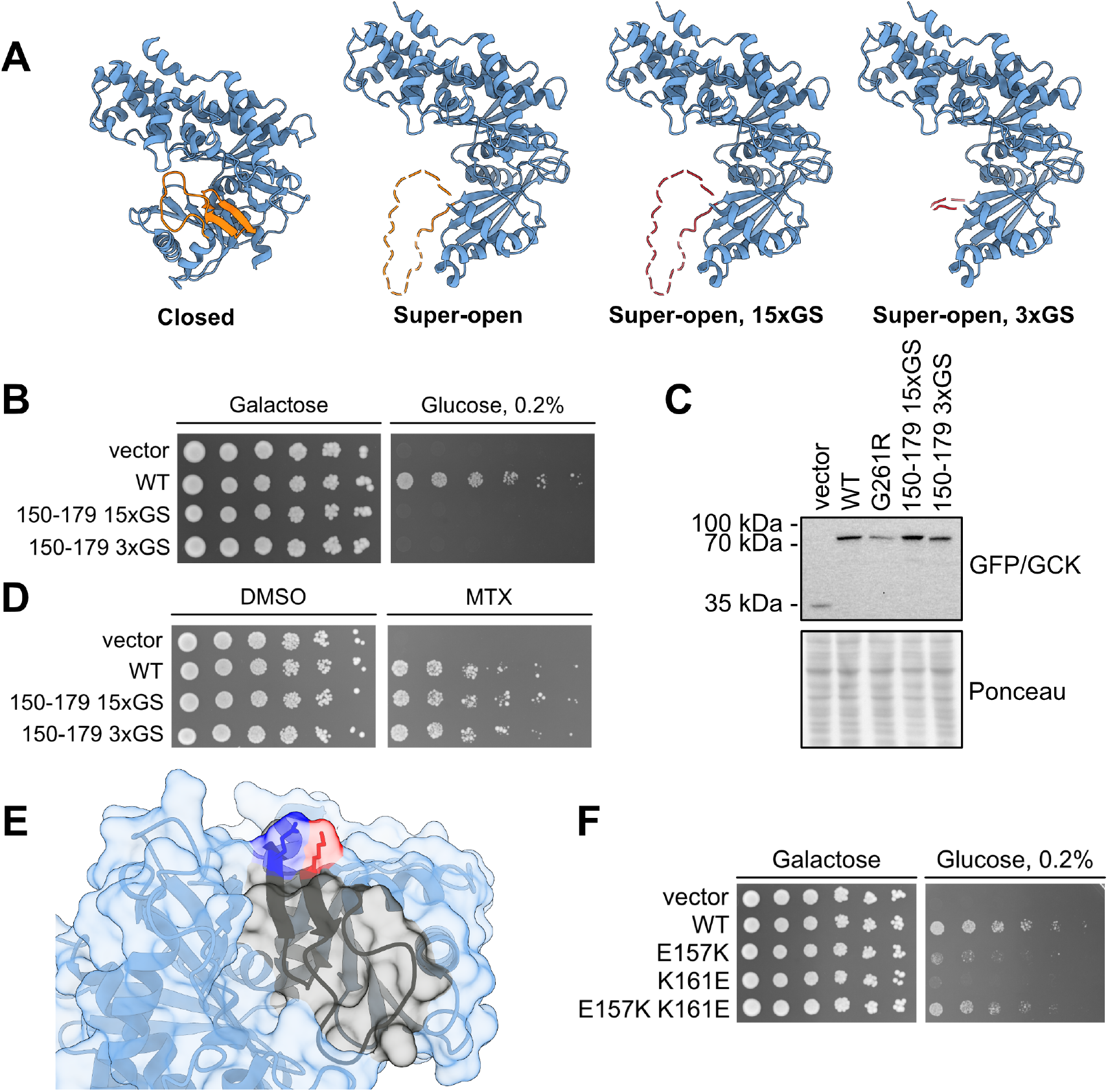
A conformational shift towards the super-open state as a mechanism for variants in the 150–179 region. (A) Left, protein structures of wild-type glucokinase in the closed and super-open states with the 150–179 marked in orange. Right, overview of glucokinase in the super-open state with the 150–179 region substituted by 30 (15xGS) or 6 residues of GS (3xGS) shown in red. (B) Yeast growth assay scoring the activity of wild-type glucokinase (WT) and the two mutants. The growth on galactose is used as a control while growth on 0.2% glucose reflects glucokinase activity. (C) Western blot showing the protein levels of the indicated constructs expressed in the *hxk1*Δ *hxk2*Δ *glk1*Δ yeast strain from panel B. (D) DHFR-PCA probing the abundance of wild-type glucokinase (WT) and the two mutants by growing yeast cells on control medium (DMSO) and medium with methotrexate (MTX) to select for abundance. (E) Structure of glucokinase in the closed state with the 150–179 region marked in black, E157 in dark blue and K161 in red. (F) Yeast growth assay scoring the activity of wild-type glucokinase (WT) and the indicated single and double mutants. PDBs: 1V4S (closed) and 1V4T (super-open).

If the super-open state is less destabilized than the closed state by mutations in the 150–179 region, variants are expected to shift the conformational equilibrium towards the super-open state, in turn resulting in decreased activity. Accordingly, re-stabilizing the closed state should increase activity. To test this, we focused on two residues in the 150–179 region, E157 and K161, that in the crystal structure of the closed state form an ion pair (Fig. 4E). Single mutants at these positions that reverse the charges, E157K and K161E, decrease activity but not abundance, based on their high-throughput assay scores (E157K activity: -0.13 abundance: 0.96; K161E activity: 0.56 abundance: 0.92). A likely explanation is a conformational shift to an inactive state due to charge repulsion in the closed state. In turn, the closed state should become favorable again when reversing both charges using the double mutant E157K K161E, leading to increased activity relative to the single mutants. When we examined the activity of the mutants, the double mutant rescued the decreased activity of the single mutants (Fig. 4F), consistent with an increased population of the closed state.

Collectively, the above experiments support that variants in the 150–179 region decrease GCK activity by shifting the conformational ensemble towards inactive states. We cannot exclude that mutations in the region may cause local unfolding without affecting the global protein conformation. However, a prior study found that the 150–179 region folded in absence of glucose when mutating the C-terminal helix 13 (***Larion et al., 2012***). As other structural elements in the small domain affect the folding of the 150–179 region, it seems reasonable that variants causing the region to unfold would affect the entire domain’s conformation.

## Conclusions

Missense variants may perturb protein function through various mechanisms. Dissecting variant mechanisms allows us to gain insights into protein function and potentially to interfere with disease-causing variants. The development of multiplexed assays of variant effects (MAVEs) (also known as deep mutational scanning (DMS)) (***Fowler and Fields, 2014***; ***Fowler et al., 2010***) has enabled us to disentangle variant mechanisms on a massive scale by probing the effects of variants using multiple read-outs (***Chiasson et al., 2020***; ***Suiter et al., 2020***; ***Cagiada et al., 2021***; ***Jepsen et al., 2020***; ***Høie et al., 2022***; ***Amorosi et al., 2021***; ***Matreyek et al., 2021***).

Building on our prior study on GCK variant activity (***Gersing et al., 2022***), we here explored GCK variant mechanisms using a multiplexed assay reporting on cellular protein abundance. Our abundance scores included 95% of the possible nonsense and missense variants. Amino acid substitutions that decreased abundance were enriched in buried residues of the large domain. For this domain, loss of abundance therefore appears to be a general mechanism for loss-of-function variants. Accordingly, we find that 43% of variants that decrease activity do so together with abundance. The remaining 57% low-activity variants may instead perturb functional sites, such as catalytic residues, allosteric residues or residues modulating GCK conformational dynamics.

Accordingly, in the dynamic small domain variants often perturbed activity but not abundance. This domain attains multiple conformations in GCK’s catalytic cycles (***Kamata et al., 2004***), and these dynamics are crucial for appropriate GCK activity and regulation. Prior studies have focused mainly on hyperactive variants that affect the conformations and dynamics of GCK (***Larion et al., 2012***; ***Heredia et al., 2006a***,b; ***Gersing et al., 2022***). For hypoactive variants, molecular dynamics simulations have predicted five variants to shift GCK into the super-open inactive state (***Zhang et al., 2006***). To further examine such a mechanism for hypoactive variants, we extended our previous predictions of changes in protein thermodynamic stability for the closed and super-open states (***Gersing et al., 2022***), and found that variants predicted to shift GCK into the inactive state are enriched in the 150–200 and 450 regions. Consistent with the molecular dynamics simulations (***Zhang et al., 2006***), we found the five variants (Y61S, I159A, A201R, V203E, V452S) to cause a relative destabilization of the closed state, potentially leading to a shift towards the inactive state. In contrast to the prior molecular dynamics simulations and kinetic studies, however, using protein stability predictions allowed us to examine the conformational shift mechanism widely.

While computational predictions allowed us to broadly examine the conformational shift mechanism, we experimentally supported our findings focusing on the 150–179 region. This region undergoes dramatic structural changes between the different GCK conformations, forming a *β*-hairpin in the closed state while being disordered in the super-open state (***Kamata et al., 2004***). The region was tolerant to mutations in our abundance assay. Accordingly, we could replace the region by a small linker sequence without perturbing GCK’s cellular protein abundance, supporting that variants in the region are tolerated due to the region’s disorder in the super-open state. Hence, variants may decrease activity by preferentially populating the super-open state due to differential destabilization of conformations. In turn, activity should increase by stabilizing the closed state. We tested this prediction using a double mutant assumed to stabilize the closed active state, and found the mutant to rescue GCK activity. Collectively, our results support that hypoactive variants may act by a relative destabilization of the closed state causing a conformational shift to the super-open inactive state.

In summary, we used a multiplexed abundance assay to identify variants that affect GCK protein stability and conformational dynamics. By identifying the mechanistic bases of hypoactive variants, we pinpointed the residues regulating stability and dynamics to ensure appropriate GCK activity. In turn, sites where such residues concentrate may be targeted to modulate GCK activity.

## Materials and Methods

### Buffers

SDS sample buffer (4×): 250 mM Tris/HCl, 40% glycerol, 8% SDS, 0.05% pyronin G, 0.05% bromophenol blue, pH 6.8. SDS sample buffer was diluted to 1.5× in water before use and 2% *β*-mercaptoethanol was added. TE buffer: 10 mM Tris/HCl, 1 mM EDTA, pH 8.0. PBS: 6.5 mM Na_2_HPO_4_, 1.5 mM KH_2_PO_4_, 137 mM NaCl, 2.7 mM KCl, pH 7.4. Wash buffer: 50 mM Tris/HCl, 150 mM NaCl, 0.01% Tween-20, pH 7.4.

### Plasmids

The DNA sequence of pancreatic human GCK (Ensembl ENST00000403799.8) was codon optimized for yeast and cloned into pDONR221 (Genscript). Selected missense variants were generated by Genscript. To generate a destination vector for the DHFR-PCA, a Gateway cassette was inserted at the C-terminus of DHFR[F3] in pGJJ045 (***Faure et al., 2022***) (Genscript). *GCK* was cloned into the pDEST-DHFR-PCA destination vector using Gateway cloning (Invitrogen). For the GCK activity assay, *GCK* was cloned into pAG416GPD-EGFP-ccdB (Addgene plasmid 14316; http://n2t.net/addgene:14316; RRID:Addgene_14316) (***Alberti et al., 2007***) using Gateway cloning (Invitrogen).

### Yeast strains

BY4741 was used as the wild-type strain. The *hxk1*Δ *hxk2*Δ *glk1*Δ strain used for the GCK yeast complementation assay was generated previously (***Gersing et al., 2022***). Wild-type yeast cells were cultured in synthetic complete (SC) medium (2% D-glucose, 0.67% yeast nitrogen base without amino acids, 0.2% drop out (USBiological), (76 mg/L uracil, 76 mg/L methionine, 380 mg/L leucine, 76 mg/L histidine, 2% agar)) and Yeast extract-Peptone-Dextrose (YPD) medium (2% D-glucose 2% tryptone, 1% yeast extract). *hxk1*Δ *hxk2*Δ *glk1*Δ yeast cells were cultured in SC and YP medium containing D-galactose instead of D-glucose. Yeast transformations were performed as described before (***Gietz and Schiestl, 2007a***).

### Yeast growth assays

For growth assays, yeast cells were grown overnight and were harvested in the exponential phase (1200 g, 5 min, RT). Cell pellets were washed in sterile water (1200 g, 5 min, RT), and were resuspended in sterile water. The cultures were adjusted to an OD_600nm_ of 0.4 and were diluted using water in a five-fold serial dilution. The cultures were spotted in drops of 5 μL onto agar plates. The plates were briefly air dried and were incubated at 30 °C (activity assay) or 37 °C (DHFR-PCA) for two to four days.

#### DHFR-PCA

To assay for GCK variant abundance, the DHFR-PCA was used (***Pelletier et al., 1999***; ***Campbell-Valois et al., 2005***; ***Levy et al., 2014***; ***Faure et al., 2022***). For plates, SC medium with leucine, methionine, and histidine was used. For selection, a final concentration of 100 μg/mL methotrexate (Sigma-Aldrich, 100 mM stock in DMSO) and 1 mM sulfanilamide (Sigma-Aldrich, 1 M stock in acetone) were used. For control plates, a corresponding volume of DMSO was used. Plates were incubated for four days at 37 °C. As a vector control for DHFR-PCAs, pAG416GPD-EGFP-ccdB was used.

#### GCK activity assay

To assay for GCK activity, yeast cells were grown on SC medium without uracil containing 0.2 % D-glucose for three days at 30 °C.

### Protein extraction

Yeast protein extraction was performed as described before (***Kushnirov, 2000***). Accordingly, 1.5– 3 OD_600nm_ units of exponential yeast cells were harvested in Eppendorf tubes (17,000 g, 1 min, RT). Proteins were extracted by shaking cells with 100 μL of 0.1 M NaOH (1400 rpm, 5 min, RT). Then, cells were spun down (17,000 g, 1 min, RT), the supernatant was removed, and pellets were dissolved in 100 μL 1.5x SDS sample buffer (1400 rpm, 5 min, RT). Samples were boiled for 5 min prior to SDS-PAGE.

### Electrophoresis and blotting

To examine GCK protein levels, proteins in yeast extracts were separated by size on 12.5% acrylamide gels by SDS-PAGE. Subsequently, proteins were transferred to 0.2 μm nitrocellulose membranes. Following western blotting, membranes were blocked in 5% fat-free milk powder, 5 mM NaN_3_ and 0.1% Tween-20 in PBS. Then, membranes were incubated overnight at 4 °C with a primary antibody diluted 1:1000. Membranes were washed 3 times 10 minutes with Wash buffer prior to and following a 1 hour incubation with a peroxidase-conjugated secondary antibody. For detection, membranes were incubated for 2–3 minutes with ECL detection reagent (Amersham GE Healthcare), and were then developed using a ChemiDoc MP Imaging System (Bio-Rad). The primary antibody was anti-GFP (Chromotek, 3H9 3h9-100). The secondary antibody was HRP-anti-rat (Invitrogen, 31470).

#### Western blot quantification

To quantify protein levels from western blots, the Image Lab Software (Bio-Rad) was used. The software was used to measure the background-adjusted intensity of protein bands and the intensity of the Ponceau stain in the same lane. Then, a loading normalization factor was calculated for all lanes by dividing the ponceau intensity of lane 1 with that of all other lanes. Band intensities were adjusted by multiplying with their corresponding loading normalization factor. Finally, the loading-adjusted variant intensities were divided by the wild-type GCK intensity to obtain a normalized intensity that could be compared between replicates.

### Glucokinase library

#### Cloning

Three regional pENTR221 libraries spanning aa 2–171 (region 1), 172–337 (region 3), and 338–465 (region 3) of the GCK sequence were previously generated (***Gersing et al., 2022***). To clone the entry libraries into the DHFR-PCA destination vector, each regional entry library was used for a large-scale Gateway LR reaction consisting of: 169.6 ng pENTR221-GCK library, 450 ng pDEST-DHFR-PCA vector, 6 μL Gateway LR Clonase II enzyme mix (ThermoFisher), TE buffer to 30 μL. The LR reactions were incubated overnight (RT). The following day, each reaction was terminated by incubation with 3 μL proteinase K (37 °C, 10 min). For each region, 4 μL LR reaction was transformed into 100 μL NEB 10-beta electrocompetent *E. coli* cells. Following electroporation, cells were recovered in NEB 10-beta outgrowth medium (37 °C, 1 hour). Then, cells were plated on LB medium with ampicillin and incubated overnight at 37 °C. If at least 500,000 colonies were obtained, cells were scraped from plates using sterile water. Plasmid DNA was extracted from cells corresponding to 400 OD_600nm_ units (Nucleobond Xtra Midiprep Kit, Macherey-Nagel).

#### Yeast transformation

To express the GCK variant libraries in yeast, each regional plasmid library was transformed into the BY4741 yeast strain as described before (***Gietz and Schiestl, 2007b***) using the 30x scale-up. Briefly, yeast cells were grown overnight at 30 °C until late exponential phase. Cultures were then diluted with 30 °C YPD medium to an OD_600nm_ of 0.3 in a minimum volume of 150 mL, and were incubated with shaking for 4-5 hours until two divisions had occurred. Then, cells were harvested and washed two times in sterile water (1200 g, 5 min, RT). The cell pellet was resuspended in a transformation mix consisting of: 7.2 mL 50% PEG, 1.08 mL 1.0 M LiAc, 300 μL 10 mg/mL single-stranded carrier DNA, 30 μg plasmid library, sterile water to 10.8 mL. The cell suspension was incubated in a 42 °C water bath for 40 minutes with mixing by inversion every 5 minutes. Cells were harvested (3000 g, 5 min, RT), the supernatant was removed, and cells were resuspended in 30 mL sterile water. To assess the transformation efficiency, 5 μL cells were plated in duplicate on SC-uracil medium. The remaining cells were diluted in SC-uracil medium to an OD_600nm_ of 0.2, and the cultures were incubated at 30 °C with shaking for two days until saturation.

If a minimum of 500,000 transformants were obtained, two cell pellets of 9 OD_600nm_ units were harvested (17,000 g, 1 min, RT) and stored at -20 °C prior to DNA extraction to serve as technical replicates of the pre-selection condition.

In parallel to the library transformations, pEXP-DHFR-PCA wild-type GCK was transformed into the BY4741 yeast strain using the small-scale transformation protocol (***Gietz and Schiestl, 2007a***).

#### Selection

To select for GCK variant abundance, the yeast libraries were grown in duplicate on medium containing 100 μg/mL methotrexate and 1 mM sulfanilamide. For each regional yeast library, 20 OD_600nm_ units of cells were harvested in duplicate and were washed three times with sterile water (1200 g, 5 min, RT). The cells were resuspended in 500 μL sterile water and each replicate was plated on a BioAssay dish (245mm × 245mm) containing SC+leucine+methionine+histidine medium with 100 μg/mL methotrexate (Sigma-Aldrich) and 1 mM sulfanilamide (Sigma-Aldrich). The plates were incubated for four days at 37 °C. Following incubation, cells were scraped off each plate using 30 mL sterile water. Cell pellets of 9 OD_600nm_ units were harvested (17,000 g, 1 min, RT) and stored at -20 °C prior to DNA extraction.

In parallel, yeast cells expressing pEXP-DHFR-PCA wild-type GCK were also used for selection as described above but using 2.6 OD_600nm_ units of yeast cells for each replicate, which were plated on petri dishes.

Plasmid DNA was extracted from yeast cells for two replicates pre- and post-selection, both for regional libraries and a wild-type GCK control. To extract plasmid DNA, the ChargeSwitch Plasmid Yeast Mini Kit (Invitrogen) was used.

#### Sequencing

In order to calculate the change in frequency of variants following selection, we sequenced the *GCK* ORF in plasmids extracted pre- and post-selection. Sequencing was done in 14 tiles spanning the *GCK* ORF, such that each regional library was covered by 4 or 5 tiles: region 1 (tile 1–5), region 2 (tile 6–10), and region 3 (tile 10–14). The short tiles enabled sequencing of both strands in each tile to reduce base-calling errors.

First, the plasmid DNA extracted from yeast cells was adjusted to equal concentrations, and was used for a PCR to amplify each tile. Each PCR consisted of: 20 μL Phusion High-Fidelity PCR Master Mix with HF Buffer (NEB), 1 μL 10 μM forward primer, 1 μL 10 μM reverse primer, 18 μL plasmid library template. The following PCR program was used: 98 °C 30 sec, 21 cycles of 98 °C 10 sec, 63 °C 30 sec, 72 °C 60 sec, followed by 72 °C 7 min and 4 °C hold. Primer sequences can be found in the supplementary data (SKG_tilenumber_fw/rev).

Following tile amplification, Illumina index adapters were added to allow for multiplexing. For each indexing PCR, the following was mixed: 20 μL Phusion High-Fidelity PCR Master Mix with HF Buffer (NEB), 2 μL 10 μM i5 indexing adapter, 2 μL 10 μM i7 indexing adapter, 1 μL 1:10 diluted PCR product, 15 μL nuclease-free water. The following PCR program was used: 98 °C 30 sec, 7 cycles of 98 °C 15 sec, 65 °C 30 sec, 72 °C 120 sec, followed by 72 °C 7 min and hold at 4 °C.

Following the indexing PCR, the indexed DNA fragments were pooled using equal volumes, and 100 μL were run on a 4% E-gel EX Agarose Gel (Invitrogen) prior to gel extraction. Then, the quality and fragment size of the extracted DNA were examined using a 2100 Bioanalyzer system (Agilent), and the DNA concentration was adjusted using Qubit (ThermoFisher), before paired-end sequencing of the libraries using an Illumina NextSeq 550.

#### Data analysis

The TileSeqMave (https://github.com/jweile/tileseqMave, version 1.1.0) and TileSeq mutation count (https://github.com/RyogaLi/tileseq_mutcount, version 0.5.9) pipelines were used to calculate variant abundance scores from sequencing data.

#### Error calculation

Standard errors of abundance scores were calculated and refined using TileSeqMave (https://github.com/jweile/tileseqMave, version 1.1.0). In this pipeline, Bayesian refinement or regularization (***Baldi and Long, 2001***) is used to obtain more robust estimates of the standard errors. Briefly, linear regression of the fitness score and read counts from the pre-selection condition are used to obtain the prior estimate of the standard error. The empirical standard error is combined with the prior using Baldi and Long’s original formula, where *σ*_0_ represents the prior estimate of the standard error, *ν*_0_ is the degrees of freedom given to the prior estimate, *n* represents the number of experimental replicates, and *s* is the empirical standard error:

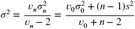

### Computational analyses

#### Defining low-abundance threshold

To set a threshold for the abundance scores, we fitted the abundance score distribution using three Gaussians. These Gaussians represent the score distributions of variants with an abundance score comparable to nonsense variants, intermediate variants, and synonymous variants, respectively. To define a cutoff for variants with decreased abundance, we used the intersection of the second and last Gaussian.

#### Structure modelling and visualisation

Protein structures were visualized and rendered using UCSF ChimeraX (v1.4), developed by the Resource for Biocomputing, Visualization, and Informatics at the University of California, San Francisco (***Pettersen et al., 2021***; ***Goddard et al., 2018***). The region spanning residues 157–179 missing from the crystal structure of the GCK super-open conformation (PDB: 1V4T) is shown in stippled lines in Fig. 4, but was modelled using Modeller (***Sali and Blundell, 1993***) to be able to obtain ΔΔ*G* estimates for variants in the region. Five structural models were generated with the Model Loops interface for Modeller available in ChimeraX (v1.3) using the super-open GCK structure (PDB: 1V4T) and the canonical GCK sequence (UniProt: P35557-1) as inputs. HETATM records and non-native terminal residues were removed from the PDB file using pdb-tools v2.4.3 (***Rodrigues et al., 2018***) prior to the loop structure generation. Model Loops was run using the standard protocol, modelling only internally missing structure, and without allowing for any remodelling of residues adjacent to the missing segment.

#### Calculation of thermodynamic stability changes

Changes in protein thermodynamic stability (ΔΔ*G* = Δ*G*_variant_ − Δ*G*_wildtype_) caused by single residue substitutions were predicted with Rosetta (GitHub sha1 c7009b3115 c22daa9efe2805d9d1ebba08-426a54) using the Cartesian ddG protocol (***Park et al., 2016***). Structure preparation and relaxation and the following ΔΔ*G* calculations were performed using an in-house pipeline (https://github.com/KULL-Centre/PRISM/tree/main/software/rosetta_ddG_pipeline, v0.2.1). Rosetta ΔΔ*G* output values were divided by 2.9 to convert from Rosetta energy units to kcal/mol (***Park et al., 2016***; ***Jepsen et al., 2020***).

ΔΔ*G* predictions for all possible point mutations in the segment spanning residues 157–179 were calculated for the super-open conformation of GCK based on the structural models created as described in the above. Predictions were performed for each of the five different structural models individually and subsequently averaged. The ΔΔ*G* predictions reported in Fig. 3 for residues 157– 179 of the super-open conformation correspond to these averages. However, all other ΔΔ*G* values presented in this work are equal to the values previously reported by ***Gersing et al***. (***2022***) and hence do not take the missing loop residues of the super-open conformation into account.

#### Calculation of weighted contact number

A weighted contact number (WCN) was calculated for every residue *i* in the crystal structure of the GCK closed conformation (PDB: 1V4S) using the expression

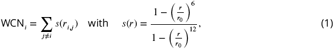

where *r*_*i,j*_ is the distance between residues *i* and *j*, and *r*_0_ = 7 Å. Interresidue distances were evaluated using the MDTraj (v1.9.7, (***McGibbon et al., 2015***)) function compute_contacts. Distances were measured as the shortest distance between any interresidue pair of atoms for residue pairs involving glycine and as the shortest distance between any two sidehchain heavy atoms for all other residue pairs.

## Acknowledgments

The authors thank Jochen Weile and Roujia Li who developed the TileSeq pipeline used for analyzing sequencing data, Vasileios Voutsinos for assistance with Illumina sequencing, Amal Al-Chaer for help with Illumina sequencing and the Bioanalyzer system, and Anne-Marie Lauridsen and Søren Lindemose for technical assistance. We acknowledge access to computing resources from the Biocomputing Core Facility at the Department of Biology, University of Copenhagen. pGJJ045 was a gift from Ben Lehner. Fig. 1A and C were created using BioRender.com.

## Data availability

Illumina sequencing data is available at the NCBI Gene Expression Omnibus (GEO) repository (accession number GSE226732). The abundance scores can be accessed from MaveDB (https://www.mavedb.org, accession number urn:mavedb:00000096-b). All data produced or analyzed in this study can be found in the supplementary data file.

## Author contributions

S.G., T.K.S., and M.C., performed the experiments. S.G., T.K.S., M.C., A.S., F.P.R., K.L.-L. and R.H.-P. analyzed the data. S.G., K.L.-L. and R.H.-P. conceived the study. S.G. wrote the paper.

