## Supplementary material for "Characterizing glucokinase variant mechanisms using a multiplexed abundance assay": Fig. S

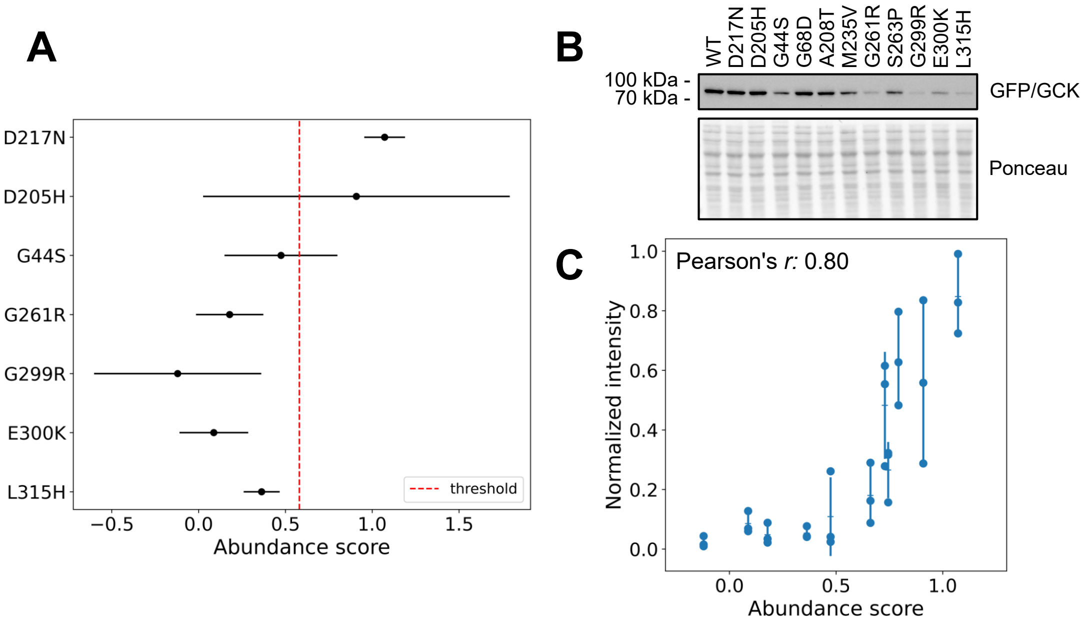

**Figure S1.** Abundance scores of test variants and correlation with western blot protein levels. (A) The abundance score and standard error of the initial test variants. The threshold for low abundance is below a score of 0.77. (B) Western blot showing the protein level of selected GCK variants spanning a wide range of abundance scores. (C) The protein levels (normalized intensity) of the 11 variants shown in panel B were quantified from three western blots. Points indicate individual western blot quantifications, horizontal lines show the mean protein level of each variant, and vertical lines show the standard deviation. The Pearson correlation between the abundance scores and three western blot quantifications is 0.80.

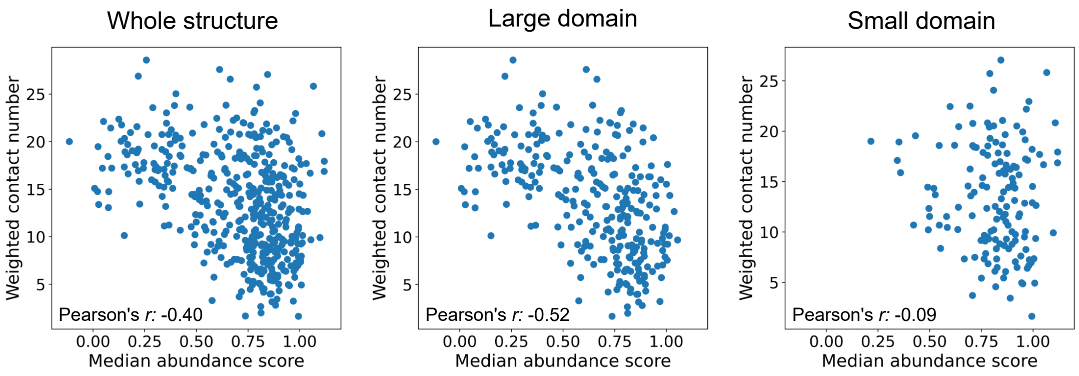

**Figure S2.** Correlation of abundance scores with weighted contact numbers. The plots show the correlation between the weighted contact number and the median abundance score for each residue. The weighted contact numbers were calculated using the closed state of GCK (PDB: 1V4S), and are a measure of residue burial, such that a higher weighted contact number indicates that a given residue is more buried. The correlation is shown for all residues in the structure and for residues in the large and small domain. For the large domain, a lower median abundance is associated with a higher contact number, suggesting that mutations at buried residues decrease abundance.

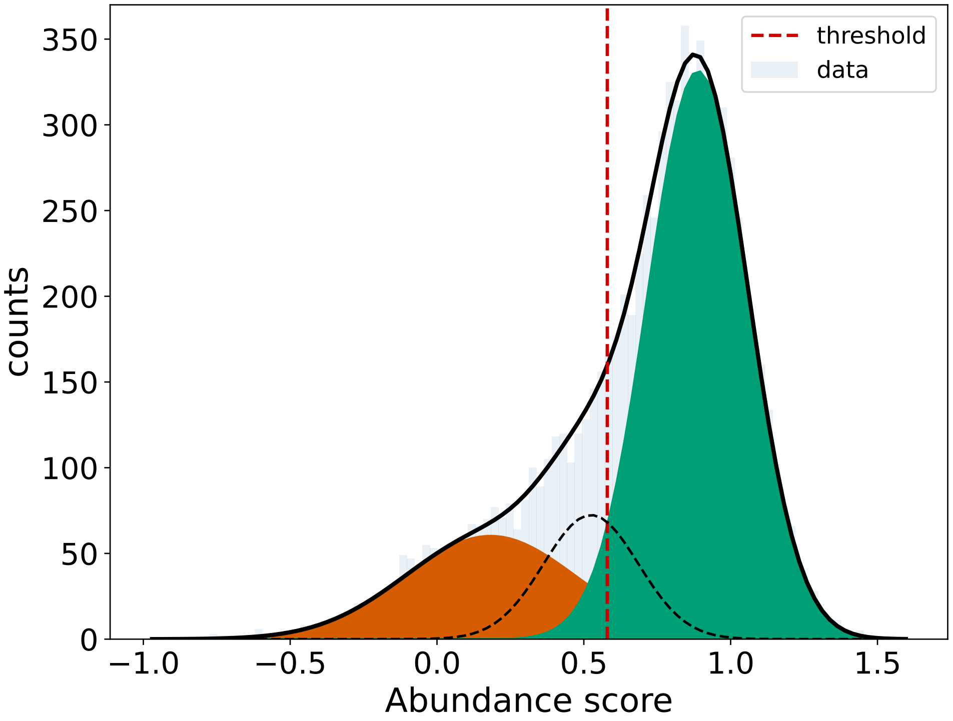

**Figure S3.** Defining a threshold for low-abundance variants. We defined a threshold for low-abundance variants by fitting the abundance score distribution (light blue) using three Gaussian distributions. We used the intersection between the second (stippled line) and last Gaussian (green) to define the cutoff (red stippled line) for variants with decreased abundance.

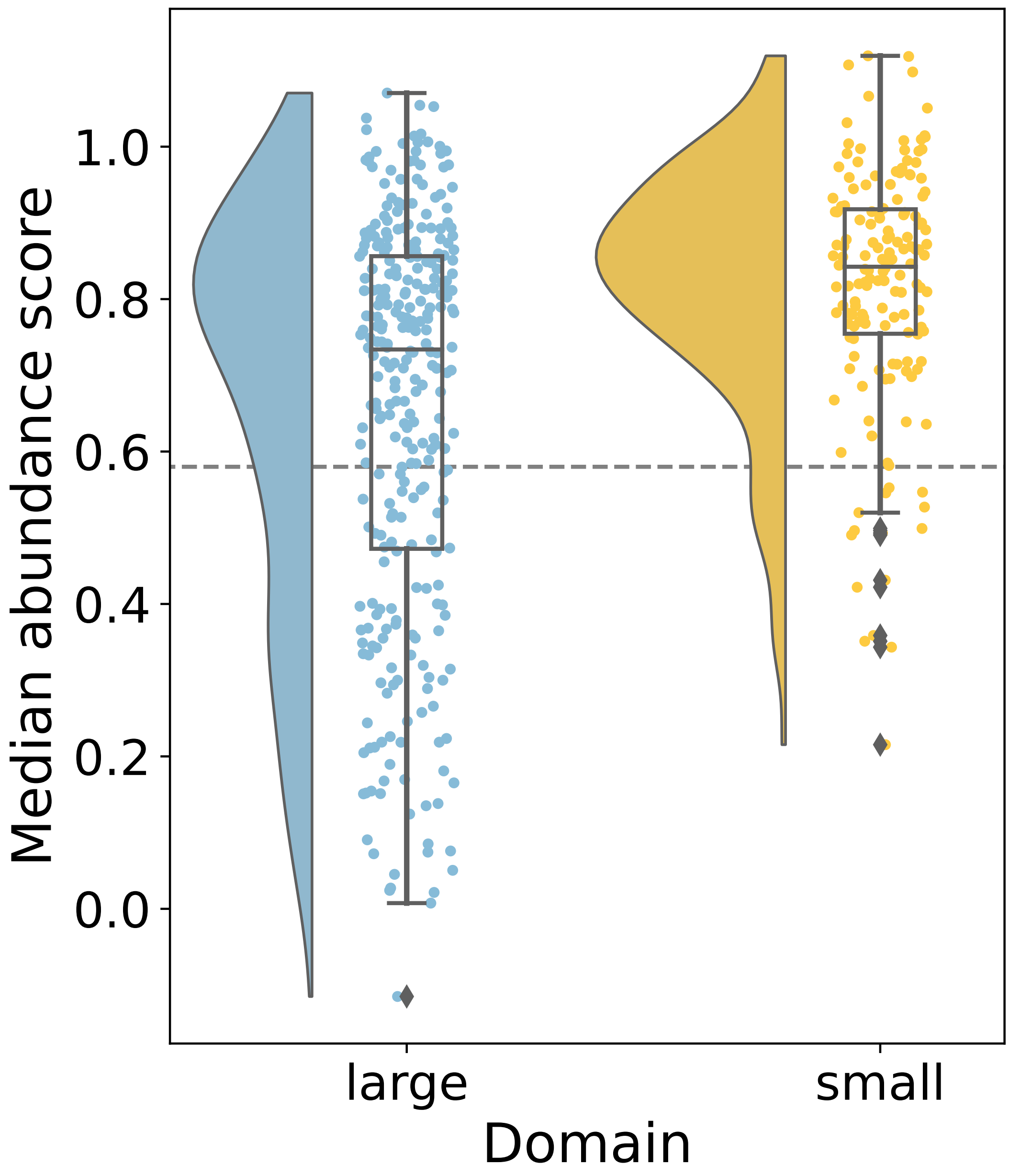

**Figure S4.** Score distributions for the two domains. The distributions of median abundance scores for residues belonging to the large or the small domain. The stippled line indicates the threshold for decreased abundance (0.58).
